# Genome-Wide Fitness Analyses of the Foodborne Pathogen *Campylobacter jejuni* in *In Vitro* and *In Vivo* Models

**DOI:** 10.1101/085720

**Authors:** Stefan P. W. de Vries, Srishti Gupta, Abiyad Baig, Elli Wright, Amy Wedley, Annette Nygaard Jensen, Lizeth LaCharme Lora, Suzanne Humphrey, Henrik Skovgård, Kareen Macleod, Elsa Pont, Dominika Wolanska, Joanna L’Heureux, Fredrick Mobegi, David Smith, Paul Everest, Aldert Zomer, Nicola Williams, Paul Wigley, Thomas Humphrey, Duncan Maskell, Andrew Grant

## Abstract

Infection by *Campylobacter* is recognised as the most common cause of foodborne bacterial illness worldwide. Faecal contamination of meat, especially chicken, during processing represents a key route of transmission to humans. There is currently no licenced vaccine and no *Campylobacter*-resistant chickens. In addition, preventative measures aimed at reducing environmental contamination and exposure of chickens to *Campylobacter jejuni* (biosecurity) have been ineffective. There is much interest in the factors/mechanisms that drive *C. jejuni* colonisation and infection of animals, and survival in the environment. It is anticipated that understanding these mechanisms will guide the development of effective intervention strategies to reduce the burden of *C. jejuni* infection. Here we present a comprehensive analysis of *C. jejuni* fitness during growth and survival within and outside hosts. A comparative analysis of transposon (Tn) gene inactivation libraries in three *C. jejuni* strains by Tn-seq demonstrated that a large proportion, 331 genes, of the *C. jejuni* genome is dedicated to (*in vitro*) growth. An extensive Tn library in *C. jejuni* M1cam (~10,000 mutants) was screened for the colonisation of commercial broiler chickens, survival in houseflies and under nutrient-rich and–poor conditions at low temperature, and infection of human gut epithelial cells. We report *C. jejuni* factors essential throughout its life cycle and we have identified genes that fulfil important roles across multiple conditions, including *maf3, fliW, fliD, pflB* and *capM*, as well as novel genes uniquely implicated in survival outside hosts. Taking a comprehensive screening approach has confirmed previous studies, that the flagella are central to the ability of *C. jejuni* to interact with its hosts. Future efforts should focus on how to exploit this knowledge to effectively control infections caused by *C. jejuni*.

**Author Summary:** *Campylobacter jejuni* is the leading bacterial cause of human diarrhoeal disease. *C. jejuni* encounters and has to overcome a wide range of “stress” conditions whilst passing through the gastrointestinal tract of humans and other animals, during processing of food products, on/in food and in the environment. We have taken a comprehensive approach to understand the basis of *C. jejuni* growth and within/outside host survival, with the aim to inform future development of intervention strategies. Using a genome-wide transposon gene inactivation approach we identified genes core to the growth of *C. jejuni*. We also determined genes that were required during the colonisation of chickens, survival in the housefly and under nutrient-rich and –poor conditions at low temperature, and during interaction with human gut epithelial tissue culture cells. This study provides a comprehensive dataset linking *C. jejuni* genes to growth and survival in models relevant to its life cycle. Genes important across multiple models were identified as well as genes only required under specific conditions. We identified that a large proportion of the *C. jejuni* genome is dedicated to growth and that the flagella fulfil a prominent role in the interaction with hosts. Our data will aid development of effective control strategies.

## Introduction

Infection by *Campylobacter* is the most common cause of foodborne bacterial diarrhoeal disease worldwide, responsible for ~96 million foodborne illnesses and ~21,000 foodborne deaths in 2010 [1]. While most cases are self-limiting, for some, campylobacteriosis is a particularly serious infection, and it is also associated with severe post-infection complications, including irritable bowel and Guillian-Barré syndromes. Consumption of undercooked poultry, unpasteurised dairy products and contaminated water represent the most common sources of human infection [2, 3]. *Campylobacter jejuni* has a broad range of environmental reservoirs that include water, birds and other domestic animals [3]. In addition, flies have been implicated as a transmission vector for *C. jejuni* for both chicken flocks and possibly also for humans as well [4-7].

*C. jejuni* encounters and has to overcome various stress conditions whilst passing through the gastrointestinal tract of humans and other animals, during processing of food products (*e.g.* slaughter process of poultry), on/in food (*e.g.* in milk or poultry meat, generally stored at low temperature) and in the environment (*e.g.* in surface water, soil, or in flies, the latter representing a transmission vector [4-7]) [8]. However, compared to other enteric pathogens such as pathogenic *Escherichia coli* and *Salmonella* spp, the survival mechanisms used by *C. jejuni* to cope with these stresses are less well-understood [9].

Based on genome analysis, the capacity of *C. jejuni* to survive outside the host and adapt to environmental stress conditions appears to be limited due to the lack of key stress regulators found in other enteric pathogens [10], however survival at low temperature in water has been reported for up to four months [9]. Establishment of colonisation and infection of host organisms by *C. jejuni* is a multifactorial process with key roles for “swimming” motility, chemotaxis, interaction with gut epithelial cells, toxin production, and adaptation to oxidative and metabolic stress [11, 12]. Factors involved in these key processes include the flagella, capsule, glycosylation systems, and two-component regulatory systems. These have all been identified as being important for chicken colonisation, human infection and environmental survival [9, 13-15].

Despite considerable research in the field, this has not led to the development and/or implementation of effective control strategies. Here, we describe the generation of extensive *C. jejuni* transposon (Tn) gene inactivation mutant libraries, and their use to comprehensively assess which genes contribute to bacterial fitness during *in vitro* growth, colonisation of commercial broiler chickens, survival in the housefly, and survival during exposure to low temperature in nutrient-rich and –poor conditions, and infection of human gut epithelial cells. This study reinforces the importance of flagella for host interactions, and identifies genes required for survival, colonisation and infection in multiple phases of the bacterium’s life cycle. Some of these may represent an Achilles’ heel of the pathogen and be a possible target for novel intervention strategies.

## Results

### Identification of Genes Required for Fitness in *C. jejuni*

To assess the genetic basis of *C. jejuni* growth and survival, genes were randomly inactivated using Tn mutagenesis in three well-characterized *C. jejuni* strains [M1cam [16, 17], NCTC 11168 (hereafter referred to as 11168) [10] and 81- 176 [18]]. Tn mutant libraries were characterised by Tn insertion site sequencing (Tn-seq [19]) (Table S1), providing a measure for the relative abundance of each Tn mutant in the library. Genes that are required for growth and survival, hereafter referred to as “fitness” genes, cannot tolerate Tn insertions, or Tn mutants in these genes are severely underrepresented in the libraries. To identify fitness genes, 23,334 unique chromosomal Tn insertions were analysed in M1cam, 15,008 in 11168, and 17,827 in 81-176 (Table S1), reaching near-saturation in terms of the number of genes that could be inactivated (Fig 1a). In addition to chromosomal Tn insertions, 2,009 and 1,919 unique insertions were in the 81-176 plasmids pVir [20] and pTet [21], respectively (Table S1). No apparent Tn insertion bias was observed (Fig S1) and each Tn library predominately yielded unique Tn insertions (Fig S2).

**Fig 1.**
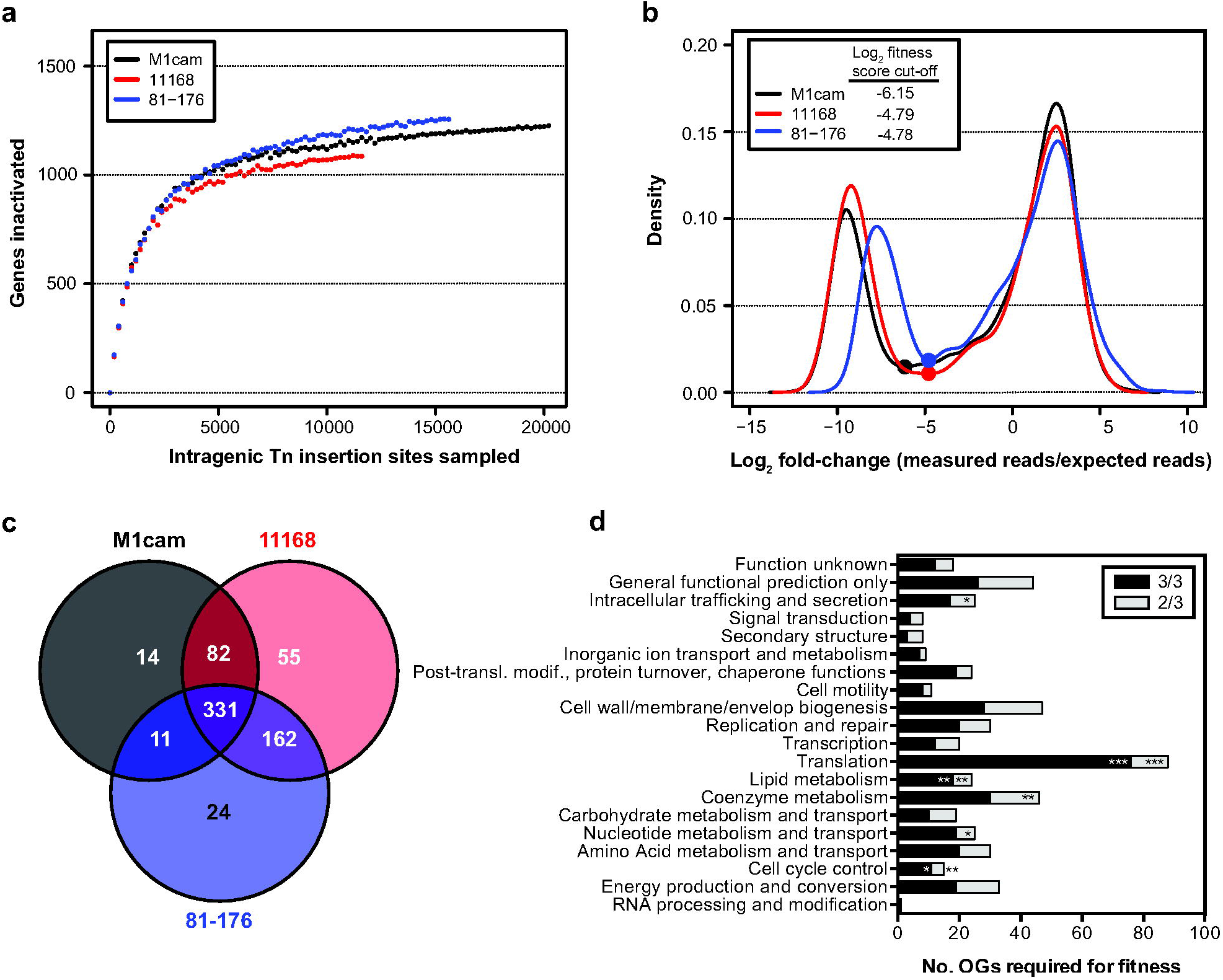
Gene fitness analysis during *in vitro* growth of *C. jejuni* M1cam, 11168 and 81-176. (A) Rarefaction analysis of intragenic Tn insertions. (B) Density plot fitness score (Log2 fold-change measured reads/expected reads) per gene. Dots indicate fitness score cut-off values. (C) Overlap of genes required for fitness in *C. jejuni* M1cam, 11168, and 81-176. (D) Functional class (COG; Cluster of Orthologous Genes) enrichment analysis of fitness genes. Fisher exact test with *Q*-value multiple testing correction; * *Q* < 0.05, ** *Q* < 0.01 and *** *Q* < 0.001.

Gene fitness score (Log2 fold-change between the observed *vs* expected of sequence reads [22]) density plots followed a bimodal distribution with the “left” population representing genes required for *in vitro* growth and survival (Fig 1b). In total, 445 genes were required for fitness in M1cam, 413 genes in 81-176 and 499 in 11168 (Table S2). Interestingly, the pTet plasmid in 81-176 harboured a single gene of unknown function (*cpp13*) that appeared to contribute to fitness. A variant of *C. jejuni* 81-176 that has lost pTet exists (D. Hendrixson, personal communication), which implies that *cpp13* is not obligate essential but may contribute to fitness, or it could be antitoxin of an uncharacterised toxin-antitoxin system. Unexpectedly, Tn insertions were observed in *dnaA* (1,323 bp) at bp position 352 in M1cam and at position 1,115 in 81-176. Consequently, *dnaA* did not pass the stringent fitness gene criteria. Disruption of *dnaA* may be tolerated at the 3’ end of the gene, in the rare event of a secondary site mutation [23], or due to the existence of merodiploids [24].

With the aim of providing a comprehensive analysis of fitness genes in the species *C. jejuni*, homologous genes were compared for the three *C. jejuni* strains (Table S2). Out of the 1,424 identified homologous gene clusters, 331 genes were required for fitness in all three strains tested and 486 genes were required in two or more strains (Fig 1c). We found that 845 gene clusters were not required for fitness in any of the analysed strains. Genes implicated in fitness were relatively dispersed across the *C. jejuni* genomes, but there were regions that were (almost) devoid of fitness genes, for example, the flagellar glycosylation gene cluster (Fig S3). The high percentage of genes required for fitness in *C. jejuni* is likely to be the consequence of its relative “minimal” genome and its proportionally large core genome [25] as well as the large number of transcriptionally coupled genes [26, 27].

The number of fitness genes shared by *C. jejuni* M1cam, 81-176 and 11168 was substantially larger, at 331, than the 175-233 genes previously reported to be obligately essential [28-31]. This may be partially due to the inclusion of genes, in our study, whose inactivation is lethal (obligate essential) as well as genes which when inactivated by a Tn result in severely compromised growth and/or survival. Further, this could be related to *C. jejuni* strain differences, growth conditions, the Tn element used, the number of Tn mutants analysed and the read-out technology. To assess the “validity” of our analysis, a systematic review of the literature on *C. jejuni* 11168 defined gene deletion and Tn mutants was performed. This analysis revealed that for 38 out of 486 (7.8%) fitness genes (required in two or more strains) identified in this study, mutants have been reported in *C. jejuni* 11168, indicating a low false-positive rate in our study. Of importance, although mutants in some of these genes could be generated, these may still result in a growth defect, *e.g.* as reported for a *C. jejuni* 11168 *pycB* mutant [32], especially if the mutant is compared against other ‘more-fit’ mutants or mutants with wild-type fitness, as is the case in the Tn screen. Our comprehensive fitness analysis indicated that a large part of the *C. jejuni* genome is dedicated to growth and survival under the condition tested, and implies that there could be opportunities for targeting some of these genes for novel intervention strategies, *e.g.* as previously reported by Mobegi *et al* [33].

Cluster of Orthologous Genes (COG) enrichment analysis showed that genes implicated in translation, lipid metabolism and cell cycle control were overrepresented amongst the genes required for fitness during *in vitro* growth/survival in all three tested *C. jejuni* strains. Extending the analysis to fitness genes required in two or more strains also showed overrepresentation of coenzyme metabolism, nucleotide metabolism and intracellular trafficking and secretion genes (Fig 1d). Required for fitness were, amongst others, genes implicated in replication, transcription, translation (40 out of 49 ribosomal protein genes), purine and pyrimidine metabolism, energy metabolism (ATP and NAD synthase, NADH-quinone oxidoreductase), isoprene biosynthesis, protein secretion (Sec and Tat pathway), as well as genes involved in cofactor biosynthesis (thiamine, folic acid and heme) and oxidative stress (see Table S2 for a complete overview). Further, the complete gluconeogenic pathway was found to be required for fitness whereas the majority of the enzymes of the tricarboxylic acid (TCA) cycle were not, except for aconitase (*acnB*), probably reflecting flexibility in this part of the bacterium’s metabolism.

*C. jejuni* expresses surface structures such as flagella, lipooligosaccharide (LOS) and capsular polysaccharides (CPS). Inactivation of genes in these pathways severely attenuated fitness. This included flagellar basal body rod proteins (encoded by *flgAC* and *fliEL*), the MS-ring (*fliF*) and C-ring (*fliG*) as well as components of the flagellar type III secretion system (*fliQH*). Genes required for formation of the LOS lipid A molecule (*lpxABCDL*), KDO (*kdsAB*) and the first L-*glycero*-D-*manno*-heptose residue (*waaC*) were required for fitness, whereas the remainder of the genes responsible for the core oligosaccharide were not. Of the capsular biosynthesis gene cluster only the inactivation of the last two genes (*kpsDF*) resulted in impaired fitness. Genes required for cell envelope generation were also important for fitness including fatty acid biosynthesis genes (*accABCD* and *fabDFGHLZ*), peptidoglycan (*dapADEF, ddl, murABCDEFG, pbpABC*), and the rod-shape determining protein genes (*mreBCD*). Protein glycosylation is tightly linked to virulence, of which the *N*-linked protein glycosylation pathway genes *pglACD* were required for fitness. This is in contrast to flagella glycosylation genes, which had no conserved role in fitness during *in vitro* growth and survival (Table S1).

### Quantitative Analysis of Genes Implicated in the Life Cycle of *C. jejuni*

The same extensive Tn library in strain M1cam (Table S1; 9,951 unique Tn insertions with 1,124 genes harbouring Tn insertions) was screened in various *in vivo* and *in vitro* models as a proxy for some of the conditions that *C. jejuni* might encounter during its life cycle from chicken-to-human. Screening the same Tn mutant library through all of the different models facilitated a comparative analysis across the models. The M1cam library was screened during the colonisation of commercial broiler chickens (natural host), survival in the housefly (transmission vector), survival under nutrient-rich- and nutrient-poor conditions at low temperature, and in models that mimic (stages of) infection of humans, *i.e.* adhesion and invasion of human gut epithelial cells. For comparative purposes we included data that we obtained in a study analysing the infection of gnotobiotic piglets (de Vries *et al.*, submitted). Tn-seq analysis after exposure to a challenge, compared with the control condition, provided a quantitative measure for the contribution of a gene to fitness in each of the models (Fig 2 and Table S3 and S4).

**Fig 2.**
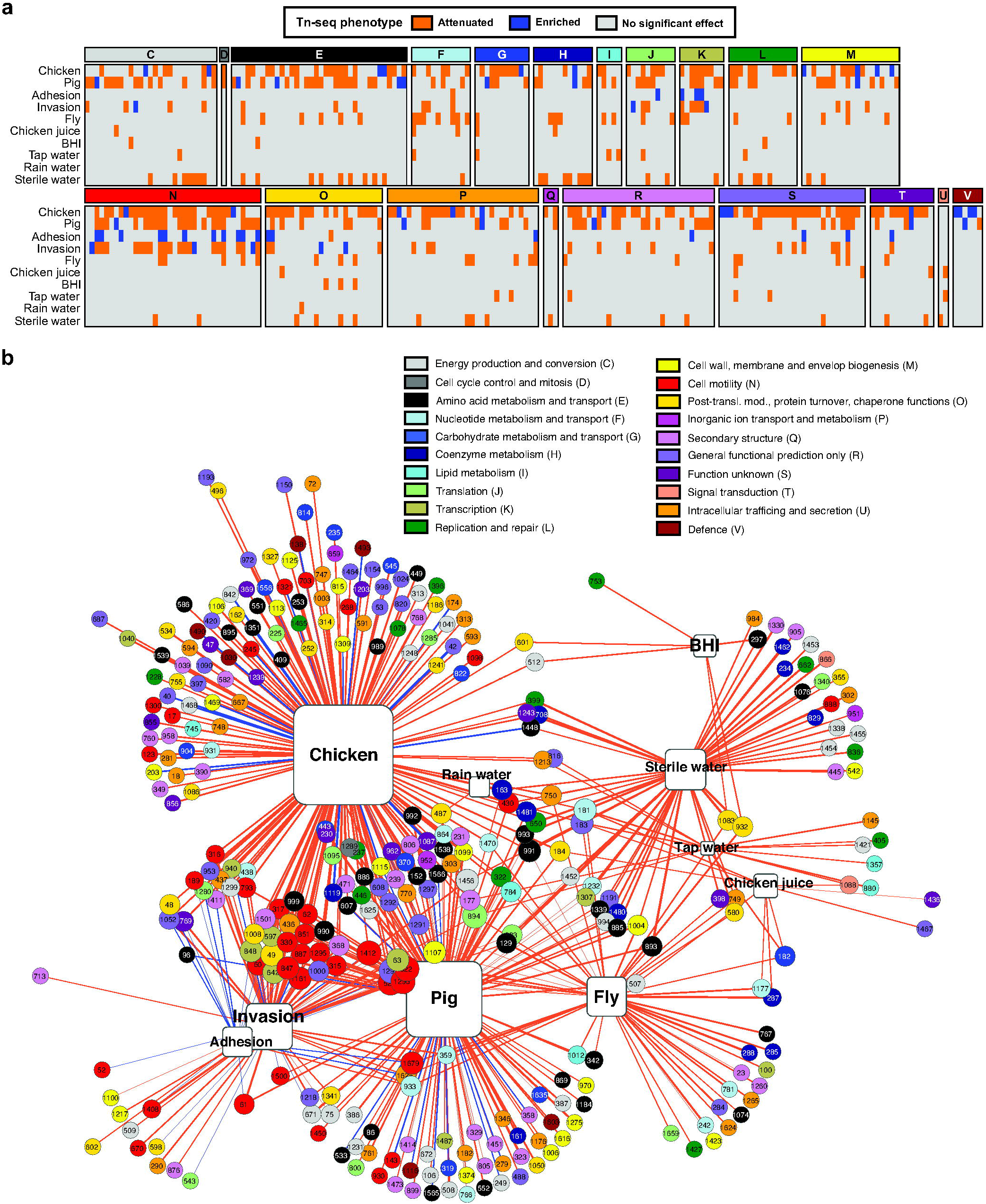
Identification of conditional essential genes in *C. jejuni*. (A) Effect of Tn insertions on the ability of *C. jejuni* M1cam to colonize commercial broiler chickens, infect gnotobiotic piglets, adhere and invade gut epithelial tissue culture cells, survive in flies and at 4°C in various media (chicken juice, BHI, tap water, rain water, and sterile water). Genes of which Tn mutants showed significantly attenuated or enriched fitness in the experimental models are shown and are grouped according to their COG functional classification. Data represented as Log2 fold-change is also presented in Table S4. (B) Gene-model interaction network showing attenuated (orange lines) and enriched (blue lines) Tn-seq scores linked to their respective models; the thickness of the connecting lines corresponds to the Log2 fold-change (input/output). The gene numbers correspond to the *C. jejuni* M1cam locus-tags [16] and are color-coded according to their COG functional class. The size of the genes and the models displayed increases with the number of interactions.

The number of Tn mutants recovered from each model confirmed that the complexity of the Tn library was maintained in survival in the housefly, under nutrient-rich and –poor conditions at 4°C and during adhesion and invasion of gut epithelial cells (Fig S4). The Tn mutant library complexity was reduced after colonisation of chickens, *i.e.* 23% of the input Tns were recovered from chickens (Fig S4), which is likely due to the existence of a population bottleneck (Fig S4). Therefore, we applied more stringent criteria for selection of candidate genes required in this model (Table S3 and Materials and Methods). For comparison, 72% of the Tn’s were recovered after infection of gnotobiotic piglets (Fig S4) (de Vries *et al.*, submitted). A detailed analysis for each of the models in this study is provided below.

### Genes Required for Colonisation of Commercial Broiler Chickens

Broilers in general become colonised with *Campylobacter* spp. at about 3-4 weeks of age [34]. We have screened the M1cam Tn library ‘C’ in a relevant chicken colonisation model, *i.e.* in 3-week-old Ross 308 commercial broiler birds. Four cages, each with 5-7 birds, were inoculated with the M1cam Tn library and 6 days post-inoculation (p.i.) colonising Tn mutants were recovered. Due to the coprophagic behaviour of chickens, each individual cage was considered a colonisation unit [35]. We hypothesised that a group-level approach would improve the robustness of the analysis and reduce any bias introduced by random dropout of Tn mutants as observed in our previous work using a signature-tagged mutagenesis approach [36] and using wild-type isogenic-tagged strains (WITS) that have indistinguishable phenotypes in pure culture [37]. Both these approaches indicated the highly complex and dynamic nature of chicken colonisation by *C. jejuni*.

On average, cages harboured 1,641 ± 226 Tn mutants (> 10 reads) compared to 7,325 ± 538 Tns in the input (Fig S4), revealing the existence of a population bottleneck, even at the cage level. However, the average read count of the four cages demonstrated recovery of 3,111 Tn mutants (> 10 reads) covering 701 genes. This indicated that, despite a population bottleneck, a large part of the genome could, with additional stringent filtering steps (see Materials and Methods), be analysed for its role during colonisation of chickens.

Tn mutants of 172 genes were significantly under-represented in colonised chickens, whereas 24 appeared to enhance fitness during colonisation (Fig 2 and Table S3 and S4). Functional class enrichment analysis revealed that genes linked to the COG class “cell motility” were significantly enriched amongst those required during colonisation (Fig 2 and Fig S5), underscoring the prominent role of motility for colonisation of chickens.

Previously, work by Johnson *et al.*, reported a Tn library screen in a 1-day-old chicken colonisation model [29]. Screening 1,155 *C. jejuni* 81-176 Tn mutants identified 130 genes that were required for colonisation and 30 genes that appeared to enhance colonisation [29]. Interpretation of the importance of the colonisation genes identified by Johnson *et al.*, is complicated due to limited validation of the Tn screen; only the importance of a single candidate gene, *mapA*, was tested and confirmed [29]. We found a limited overlap of 23 chicken colonisation genes between the two studies, 11 of which were linked to motility and the flagellar system. The datasets had one gene (CJM1_0420; hypothetical protein) in common that was beneficial for colonisation. Differences between identified colonisation genes are likely the result of the experimental model employed, *e.g.* older birds harbouring a more mature intestinal microbial community and younger birds being more permissive for colonisation by *C. jejuni, e.g.* as reflected in a lower inoculation dose to establish colonisation [38].

For validation of the chicken colonisation Tn library screen, defined deletion mutants in 19 genes were tested individually for their ability to colonise chickens. The importance in chicken colonisation was confirmed for genes involved in chemotaxis (*mcp4_1*), the flagellar system (*pflB*, *fliD, fliW*, and *maf3*), *N-*linked protein glycosylation system (*capM*, also referred to as *pglH*), and phosphate transport (*pstA*) (Fig 3a). Although motility is considered a driving factor in chicken colonisation, we have found that mutants in *maf3*, *capM* and *pstA*, that have a defect in colonisation, displayed wild-type motility (Fig S6). In the Tn library screen, mutants in *eptA* (also referred to as *eptC* [39]), *glnP*, *jlpA*, *fdhA* and CJM1cam_1125 (uncharacterised glycosyltransferase) showed reduced colonisation. However, validation with defined deletion mutants revealed slightly higher bacterial loads in chickens when compared to the wild-type (Fig 3a); motility of these defined gene deletion mutants did not differ significantly from the wild-type (Fig S6). In contrast to our findings, previous work by Cullen *et al.*, reported a slight reduction in motility after deletion of *eptC* in *C. jejuni* 81-176. EptC catalyses the phosphoethanolamine modification of FlgG and lipid A, [39], suggesting that the importance of *eptA* could be strain dependent. No difference to the wild-type was found in the colonisation proficiency of defined mutants in *moaA*, CJM1cam_0303 (hypothetical protein), CJM1cam_0438 (hypothetical protein), *flaG*, *ilvB*, *engD* and *gltA*. Identification of these genes as attenuated in the Tn library screen might be attributed to the competition effect of Tn mutants in other genes being present during the Tn library screen, whilst validation experiments were conducted with single mutant inocula. Here we opted for validation experiments with single mutant inocula due to the unpredictable outcome of competition experiments, even in the simplest case of two phenotypically indistinguishable (in pure culture) WITS [37]. It is possible that some candidate colonisation genes represent false positives due to the random loss of mutants, *i.e.* an infection bottleneck that was observed in the chicken model.

**Fig 3.**
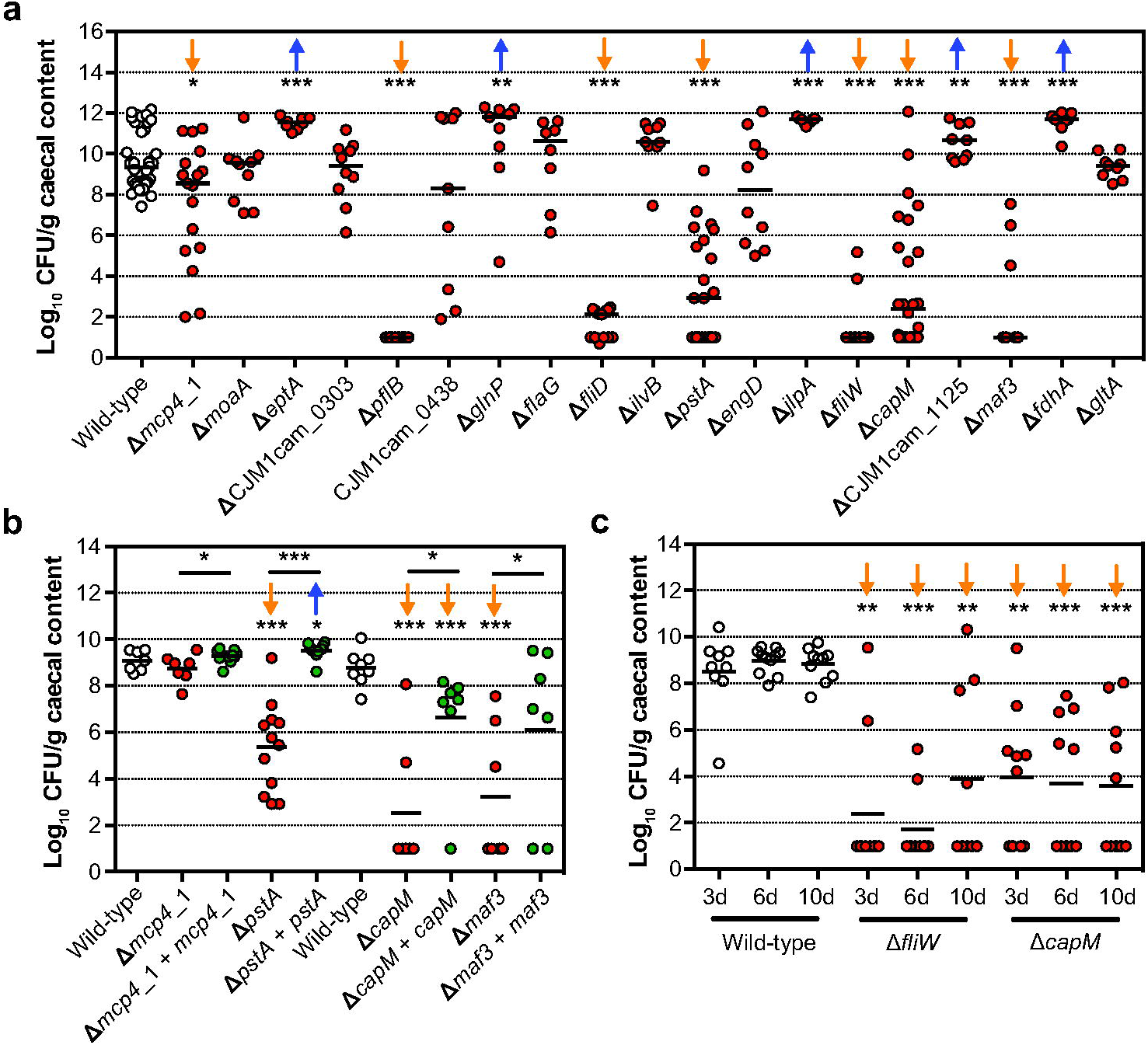
Validation of chicken colonisation Tn mutant library screen. (A) Colonisation levels of *C. jejuni* M1cam gene deletion mutants 6 d pi. (*n* ≥ 7) (B) Colonisation of gene deletion mutants and complemented gene deletion mutants (*n* ≥ 7). (C) Colonisation kinetics of *C. jejuni* M1cam wild-type and defined mutants in *fliW* and *capM* (*n* ≥ 9). Chickens being colonised *vs* not colonised were 2/8, 2/8 and 3/6 ≥ for the *fliW* mutant and 4/5, 4/6 and 5/5 for the *capM* mutant at day 3, 6 and 10 p.i., respectively. Statistical significance was analysed using a Mann-Whitney test with * *P* < 0.05, ** *P* < 0.01 and *** *P* < 0.001. The orange arrows represent an attenuated phenotype while the blue arrows represent an enriched chicken colonisation phenotype.

Genetic complementation of mutants in *pstA*, *capM* and *maf3* did significantly increase their colonisation capacity compared to their respective gene deletion mutants (Fig 3b), confirming their role in chicken colonisation. Deletion of *fliD*, encoding the flagellar cap protein, resulted in severely attenuated chicken colonisation (Fig 3a). However, attempts to complement the *fliD* deletion failed (see Fig S8 for details). When analysing colonisation of the genetically complemented *mcp4_1* mutant in chickens, we found that, in contrast to initial validation experiments (Fig 3a), the *mcp4_1* mutant was able to colonise chickens; which could be the result of intrinsically lower colonisation resistance in the batch of chickens used in this experiment.

Despite the existence of a colonisation bottleneck and the incomplete reproducibility when comparing defined deletion mutants with the results obtained in the Tn library screen, this work has identified *pstA, capM, maf3* and *mcp4*_1 as novel colonisation factors in 3-week-old broiler chickens. Previously, phosphate transport system (*pstACS*) genes in *C. jejuni* 81-176 were shown to be expressed at higher levels during colonisation of 1-day-old chickens compared to *in vitro* grown bacteria [40], which can likely be attributed to low levels of phosphate in the chicken gut. Deletion of the *N*-linked protein glycosylation gene *capM* in *C. jejuni* 81116 was previously reported to reduce colonisation of chickens [41, 42]. The genomic location of *maf3* (motility accessory factor) suggests a role in flagellar glycosylation [43]. In addition to the role of *mcp4*_1 in the colonisation of chickens identified in our study, deletion of other methyl-accepting chemotaxis proteins (Mcp) also resulted in reduced colonisation by *C. jejuni* of mice [44].

In validation experiments (Fig 3a) we observed an ‘all or nothing’ colonisation effect in some chickens infected with particular defined mutants, *e.g. pflA*, *fliD* and *fliW*, whereas for other deletion mutants, such as *capM* and *mcp4_1*, colonisation levels varied considerably between the birds. This indicates that there might be differences in the colonisation permissiveness/resistance between birds, within and between experiments. To investigate this in more detail, the colonisation of the wild-type, *fliW* and *capM* mutants were measured at different time intervals (3, 6 and 10 days p.i.) Fig 3c). Chickens were colonised at high levels from 3 days onwards and neither an increase in the levels of colonisation nor the number of colonised birds was observed. Surprisingly, the level of colonisation of the *fliW* and *capM* mutants did not increase between 3 and 10 days p.i. and there was also no obvious increase in the number of colonised chickens (Fig 3c). We hypothesise that the colonisation responses observed in our validation experiments were potentially confounded by variation in gut microbiota composition or differential inflammatory responses elicited during colonisation [40, 45, 46].

### Genes Required for Survival in the Housefly

As a transmission vector model for *C. jejuni* [4-7], survival of the M1cam Tn library ‘C’ was examined 4 hours after individual inoculation of houseflies *via* their proboscis. Tn mutants in 48 genes showed reduced survival and no genes were identified for enhanced survival (Fig 2 and Table S3 and S4). Genes of the COG class “nucleotide transport and metabolism” were overrepresented amongst the genes linked to survival in the housefly (Fig 2 and Fig S5). These were the non-essential genes in the purine (*purLMN*) and pyrimide (*pyrC*_1/2 and *pyrDF*) biosynthesis pathways. Although the Tn library screen identified a number of candidate survival genes, validation experiments with 7 defined gene deletion mutants (inoculated as single mutants) were non-confirmative (Fig 4a). Our inability to confirm the role of identified candidate genes might be the result of low levels of attenuation, or be due to the lack of competition with other mutants. However, the attenuated survival of a *capM* deletion mutant approached significance (*P* = 0.0513, two-tailed Mann-Whitney) when compared to the wild-type. As the importance of *capM* in chicken colonisation was confirmed *via* genetic complementation of the deletion mutant, we also tested the *capM* mutant alongside its genetically complemented mutant for survival in the housefly. This confirmed that *capM* is also involved in survival in the housefly (Fig 4b).

**Fig 4.**
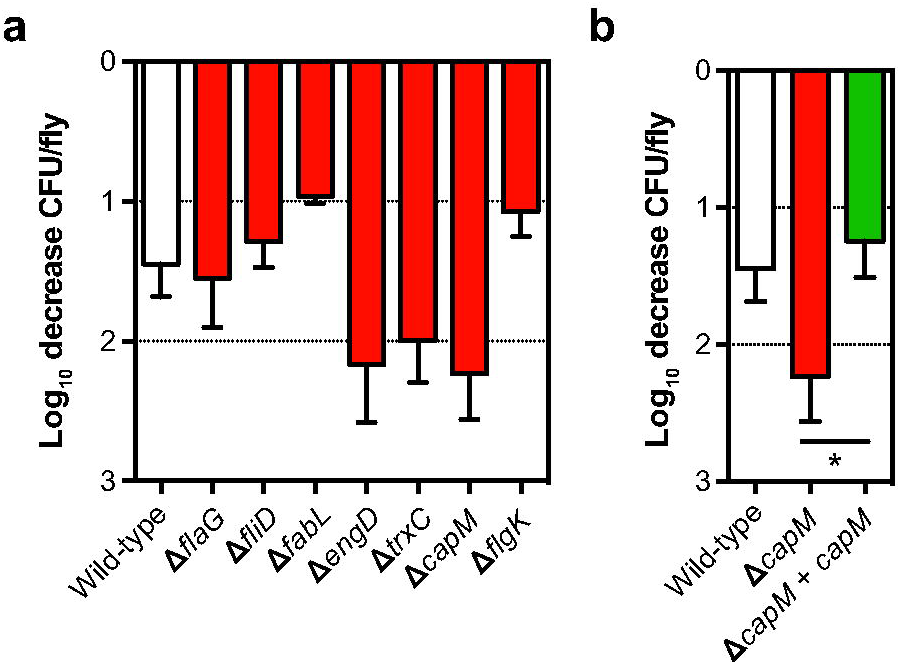
Validation of Tn mutant library screen during survival in the housefly. (A) Survival (Log10 decrease inoculum *vs* recovered) of defined *C. jejuni* M1 gene deletion mutant in the housefly after 4 h. (B) Survival of wild-type, *capM* defined gene deletion mutant and genetically complemented mutant. Data shown are Log10 decrease of CFU per fly relative to the inoculum and plotted as means with SEM (*n* ≥ 4). Significance was analysed using a Mann-Whitney test with * *P* < 0.05.

### Survival in Nutrient-Rich and –Poor Conditions at Low Temperature

To identify *C. jejuni* genes involved in survival in nutrient-rich and –poor conditions at low temperature (4°C), the M1cam Tn library ‘C’ was incubated in “chicken juice” (liquid obtained after thawing chicken carcasses), BHI broth and in water (sterile, tap and rain), with the “water” models being included to assess the survival under more general environmental conditions. The Tn mutants that survived were recovered after 7 days incubation and compared to the Tn library composition at T = 0 days. Ten genes were implicated in survival in chicken juice and 6 genes in BHI medium. Considerable variation was found for the number of genes implicated in survival at 4°C in nutrient-poor conditions, with 13 genes identified in tap water and 57 in sterile water. However, only one candidate gene, encoding the heat shock response protein ClpB, was identified as being required for survival in rain water (Fig 2 and Table S3 and S4). We suspect that these variations could arise due to differences in chemical composition, pH and potentially other bacteria present in water samples [47, 48]. Previous work in *C. jejuni* 11168 showed that *clpB* is involved in regulating a stress response to acidic pH while transiting the human stomach [49]. In addition, ClpB could facilitate the solubilisation and renaturing of aggregated proteins at low temperature under which translation is repressed [50]. This could be translated to our findings that ClpB may act as a stress response regulator and aid in the survival of *C. jejuni* under conditions of stress. Given the relatively low number of genes associated with survival at low temperature and the relatively mild attenuation, as expressed by Tn-seq fitness score, we hypothesise that survival may be a passive rather than active mechanism [51].

At low temperature, an oxidative stress response is induced in *C. jejuni* [9, 52]. In our Tn library screen, the gene encoding the oxidoreductase TrxC was found to be required for survival in sterile water and BHI at low temperature. In addition, the regulator of oxidative stress, PerR, was also linked to survival in sterile water. PerR plays a role in controlling oxidative stress resistance and survival under aerobic conditions [52, 53]. The RacRS two-component system is important for chicken colonisation and is part of a temperature-dependent signalling pathway [54]. In our study, Tn mutants in *racS* had reduced survival in chicken juice and tap water at low temperature. Chemotaxis has also been suggested to play a role in survival at low temperature [55]. In line with this, Tn mutants of *mcp4_2* were attenuated for survival in chicken juice and tap water. In addition, Tn mutants in *kefB* and *czcD* (antiporters), *fabI* (fatty acid metabolism) and CJM1cam_0181 to CJM1cam_0183 (*purN, nnr*, and a gene encoding a hypothetical protein) were linked to survival in both chicken juice and tap water.

For validation of the Tn-seq screen, 10 defined gene deletion mutants were assayed under all conditions described above. Interestingly, survival of the wild-type strain was significantly lower in chicken juice compared to any of the other conditions analysed (Mann-Whitney test, *P* < 0.001). We found that several deletion mutants were attenuated for survival after 3 days in BHI and water (sterile, tap and rain). However, at 7 days their survival did not differ significantly to that of the wild-type. This is most likely due to further decreasing levels of the wild-type between day 3 and 7 (Fig 5a).

**Fig 5.**
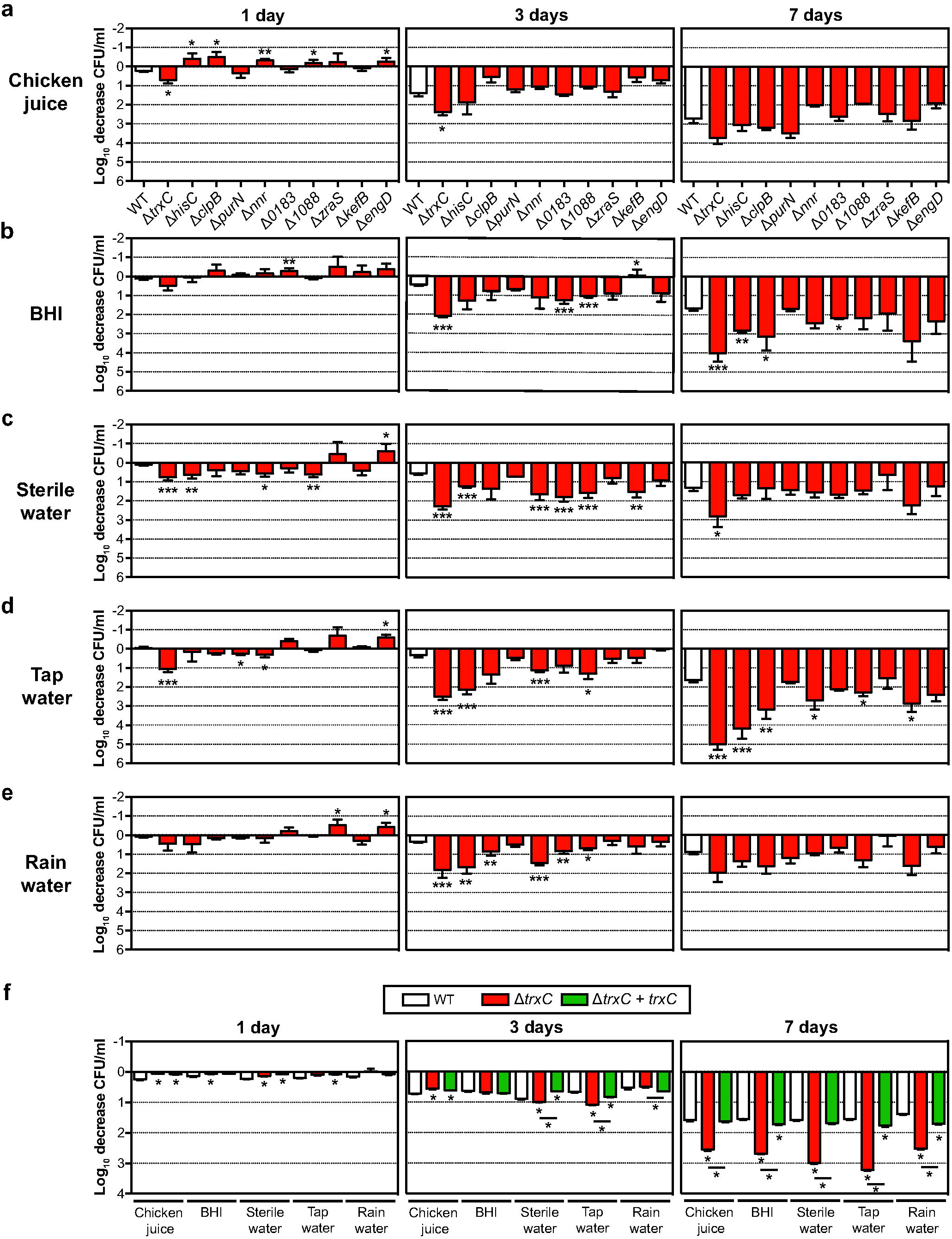
Validation of Tn mutant library screen during survival at low temperature under nutrient-rich and –poor conditions. (A-E) Survival of defined *C. jejuni* M1cam gene deletion mutant in (A) BHI, (B) chicken juice, (C) sterile water, (D) tap water, (E) rain water. Survival of *trxC* defined gene deletion mutant and complemented mutant (F), at 4°C in different media. Data shown are Log10 decrease of CFU/ml relative to the 0 day time-point and plotted as means with SEM (*n* ≥ 4). Statistical significance was analysed using a Mann-Whitney test with * *P* < 0.05, ** *P* < 0.01 and *** *P* < 0.001.

Confirmatory experiments with deletion mutants highlighted the contribution of *hisC* (aromatic amino acid aminotransferase) and *trxC* (thiol-disulphide oxidoreductase) for survival at low temperature (Fig 5b-e). The *hisC* mutant showed attenuated survival in BHI (7 days), sterile water (3 days), tap water (3 and 7 days) and rain water (3 days) (Fig 5b-e). Deletion of *trxC*, however, resulted in reduced survival of *C. jejuni* in BHI, sterile water and tap water after 7 days (Fig 5b-d) and in chicken juice and rain water after 3 days (Fig 5a,e). Although the deletion of *trxC* resulted in a prolonged lag-phase during growth *in vitro* in BHI broth (Fig S7), genetic complementation of the *trxC* deletion restored its phenotype under all conditions tested, confirming the importance of *trxC* in the survival of *C. jejuni* at 4°C, both in nutrient-rich and –poor conditions (Fig 5b). Thus far, the function of TrxC in *C. jejuni* is unknown. The closest ortholog is TrxC in *Helicobacter pylori*, which is required for protection against oxidative stress [56]. We found that the sensitivity of the M1cam *trxC* mutant to hydrogen peroxide was increased, and that sensitivity was restored to near wild type levels in the genetically complemented mutant (Fig 6). Therefore, our data support a role for *C. jejuni* TrxC in an oxidative stress response, and could be a good target for intervention strategies.

**Fig 6.**
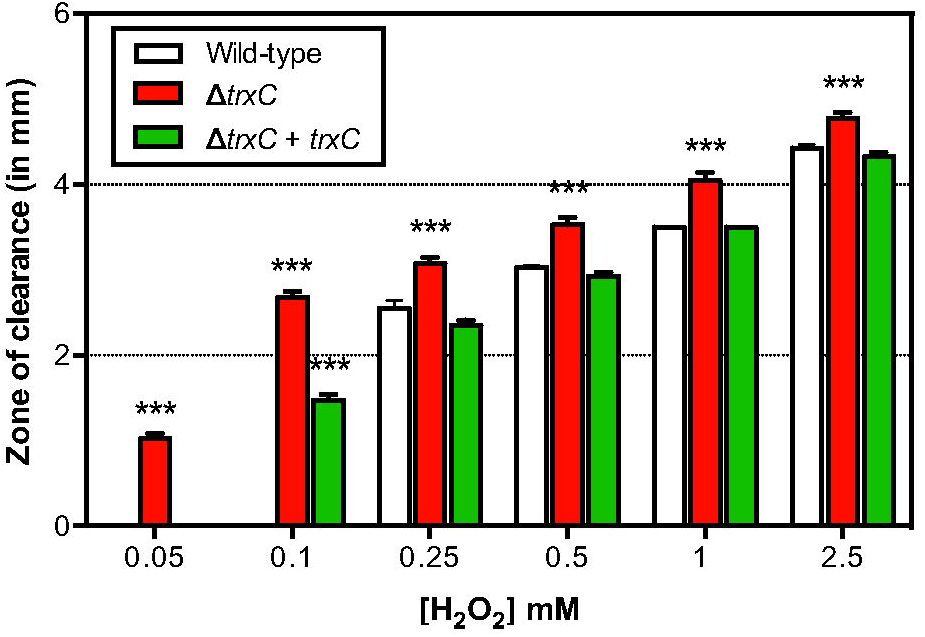
Sensitivity to hydrogen peroxide of *C. jejuni* M1cam wild-type, *trxC* defined gene deletion mutant and *trxC* complemented mutant. Data shown are zones of inhibition and plotted as means with SEM (*n* ≥ 4). Statistical significance was analysed using a two-way ANOVA test with *** *P* < 0.001.

### Adhesion and Invasion of Human Gut Epithelial Cells

The M1cam mutant library ‘C’ was used to infect Caco-2 human gut epithelial cells. To identify genes required for adhesion, Tn mutants that adhered to Caco-2 epithelial cells were compared to the non-adherent fraction rather than to the inoculum. This was to compensate for the survival of the Tn mutant library in the infection medium, *i.e.* to restrict the number of false positives due to attenuated survival in the infection medium. Comparing the Tn mutants that invaded the epithelial cells with the non-adherent fraction identified 57 candidate genes involved in cellular invasion, whereas only two genes passed our filtering criteria for adhesion (Fig 2 and Table S3 and S4).

As anticipated, genes linked to the COG class “cell motility” were significantly overrepresented amongst the genes required for invasion of Caco-2 cells (Fig 2 and Fig S5). Previous work has shown that flagellar motility plays an important role in the pathogenesis of *C. jejuni* and is considered a driving factor for host interaction [3, 28, 57, 58]. Employing a Cos-1 monkey kidney fibroblast cells invasion model, Gao *et al.*, screened a Tn mutant library in *C. jejuni* 81-176, which led to the identification of 36 invasion genes [28], of which 19 genes were also identified in this study using Caco-2 human gut epithelial tissue culture cells. A total of 38 genes were only required for invasion by M1cam and 17 only in 81-176 [28]. Of the M1cam unique invasion genes, 14 were linked to the flagellar system, including *fliK*, *flaG*, *fliD*, *fliW*, and *maf3*. The differences in invasion requirements are likely to be caused by the different cell-types used and might vary between different strains of *C. jejuni.*

Validation of invasion genes identified by Tn-seq was performed using a panel of 15 defined gene deletion mutants, this included genes not previously reported to be linked to motility and the flagellar system (Fig 7a,b). In these experiments, the role of 13 out of 15 selected genes in invasion was confirmed (Fig 7b). Although Tn-seq only detected two genes of which Tn mutants showed reduced adhesion, 14 out of 15 deletion mutants selected for confirmatory experiments displayed attenuated adhesion (Fig 7a). This apparent discrepancy might be due to differences in the experimental set-up between the Tn library screen and the validation experiments. In the Tn library screen, interactions between adhesion-deficient and -proficient Tn mutants may have compensated for adhesion of otherwise adhesion-deficient Tn mutants. However, it is also well recognised that competition between Tn mutants is a confounding factor in Tn library screens [29, 58]. This is in contrast to the validation experiments presented here, in which defined gene deletion mutants were allowed to interact with Caco-2 cells as single mutant inocula.

**Fig 7.**
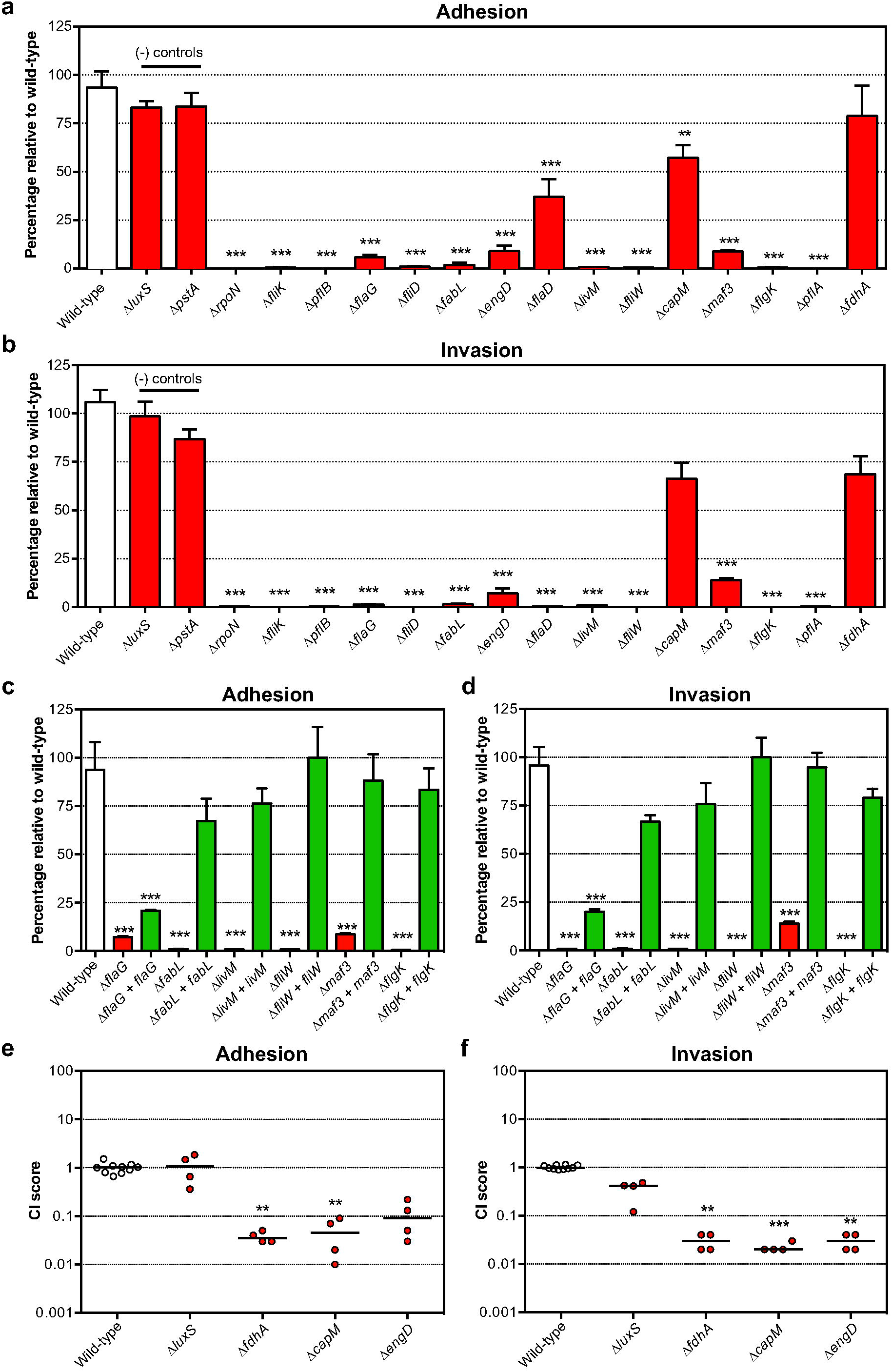
Validation of Tn mutant library screen during adhesion and invasion of human gut epithelial tissue culture cells. Adhesion to (A), and invasion (B) of, Caco-2 cells by *C. jejuni* M1cam defined gene deletion mutants. Mutants in *luxS* and *pstA*, which were not identified in our Tn-seq screen with Caco-2 cells, were included as negative controls and *rpoN*, which was previously shown to be required for adhesion and invasion [59, 60], served as a positive control. Caco-2 adhesion (C), and invasion (D), of gene deletion and genetically complemented mutants. Data is represented as percentage of wild-type (*n* ≥ 3) and plotted as means and SEM. The competitive index (CI) was calculated by dividing the ratio of mutant to wild-type bacteria recovered upon (E) adhesion to, and (F) invasion of, Caco-2 cells by the ratio of mutant to wild-type bacteria that were used in the inoculum. Statistical significance was calculated using a Mann-Whitney test where * *P* < 0.05, ** P <0.01 and *** *P* < 0.001.

Amongst the genes that were confirmed to play a role in interacting with Caco-2 cells were *livM* (amino acid metabolism), *fabL* (fatty acid metabolism) and *engD* (GTP-dependent nucleic acid-binding protein) that have no known link to the flagellar system. We assessed the motility of these mutants in semi-solid agar (Fig S6) and found that deletion of *livM* resulted in ~50% reduced motility compared to the wild-type. WGS analysis did not reveal any genomic variations in the *livM* mutant linked to motility (Table S5). The role of *engD* in colonisation of chickens was not confirmed in validation experiments with a defined deletion mutant (Fig 3). However, we confirmed the contribution of *engD* in adhesion and invasion of Caco-2 tissue culture cells (Fig 7). The *maf3* gene deletion mutant was motile (Fig S6) but had a reduced capacity to colonise chickens (Fig 3a,b) and also lacked the ability to adhere to, and invade, Caco-2 cells (Fig 7a-d).

Genetic complementation of the defined gene deletions in *fabL*, *livM*, *fliW*, *maf3* and *flgK* mutants restored their adhesion and invasion capacity to wild type-levels, confirming their role in the interaction with human gut epithelial cells (Fig 7c,d). Genetic complementation of the *flaG* deletion mutant significantly enhanced adhesion and invasion levels, however these levels were still lower than the wild-type. Although not confirmed, this may be due to deregulated *fliD* or *fliS* expression as observed for the *fliD* mutant (see Fig S8 for details). Genetic complementation of the *engD* deletion mutant was also unsuccessful due to the lack of expression of *engD* (Fig S8). Further, the *capM* and *fdhA* Tn-seq phenotypes were confirmed when mutants were assayed in competition with the wild-type (Fig. 7ef), however in mono-infection no phenotypes were observed (Fig. 7a-d). The *luxS* negative control mutant was not attenuated for adhesion and invasion in competition with the wild-type (Fig. 7ef).

## Discussion

This study presents a comprehensive analyses of gene fitness in *C. jejuni* and underlines various molecular mechanisms that are critical in its life cycle. Profiling the genes required for *in vitro* growth of three *C. jejuni* strains underscored that a large part of the genome (~27%) is vital to bacterial fitness (Table S2). Most likely as a consequence of a low redundancy in the relatively minimal genome of *C. jejuni* and transcriptional coupling of genes, as demonstrated by Dugar *et al.*, and Porcelli *et al.*, [26, 27].

Elements of the flagellar system, *i.e.* the flagellar base and periplasmic rod structure and T3SS (Fig 8), encoded by indispensable genes or genes which when inactivated by a Tn have a severe impact on fitness, provide a potential focus for intervention along with genes required for the LOS lipid A and KDO moieties. We also found that inclusion of the first L-*glycero*-D-*manno*-heptose residue (catalyzed by WaaC) was required for fitness. These structural elements have a critical role during host interaction and therefore represent promising targets for developing intervention strategies [57, 61]. Our gene fitness analysis revealed that components of the gluconeogenesis pathway were essential for the growth of *C. jejuni*. The gluconeogenic pathway is required for biosynthesis of glucose-(derivatives) that serve as building blocks for both LOS and CPS as well as the *N*- and *O*-linked protein glycosylation pathways [14]. In addition, genes (*fabZ, fabF, fabH, fabD* and *fabG*) that are part of the type II fatty acid synthesis pathway (FASII) were required for fitness in all three *C. jejuni* strains. There are several classes of FASII pathway inhibitors that have potent antimicrobial properties and consequently fatty acid synthesis is also considered a lucrative target for antibiotics [33, 62].

**Fig 8.**
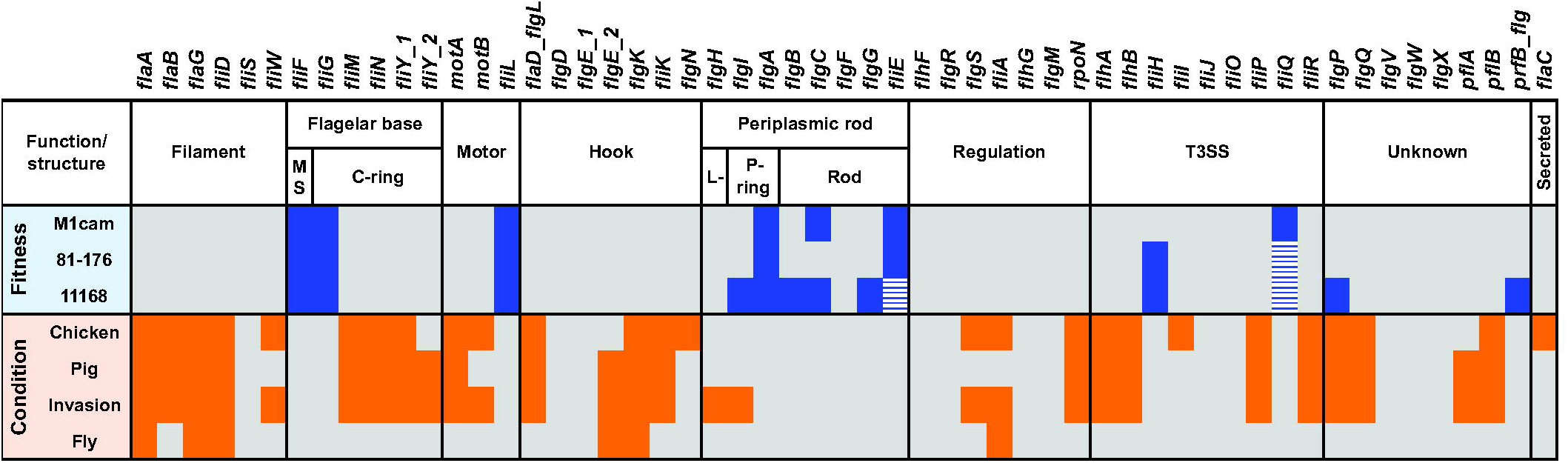
Overview of flagellar system genes required for fitness during *in vitro* growth and in model representing host interaction. Genes involved in the flagellar system that were required for fitness during *in vitro* growth of *C. jejuni* strains M1cam, 81-176 and 11168 are indicated in blue. Genes that passed the fitness score cut-off but did not pass the 0.95 probability for Tn inactivation are indicated with blue dashed boxes. Genes which in the Tn library screens were shown to be required for chicken colonisation, gnotobiotic piglet infection, invasion of human gut epithelial tissue culture cells or survival in houseflies are indicated in orange.

The same *C. jejuni* M1cam Tn library was screened in all of the experimental models presented in this study and at the same time compared with data derived from screening the same Tn library during infection of gnotobiotic piglets (De Vries *et al.*, submitted). This enabled comparative analyses of genes required in *in vitro* and *in vivo* experimental models relevant to the life cycle of *C. jejuni*. Comparing genes required for colonisation of chickens and infection of gnotobiotic piglets as well as the genes required for interaction with human gut epithelial cells, not only enabled us to identify genes associated with propagation and survival across different host species but also of genes specific to different hosts.

Twenty-eight genes were found to be required in all three host interaction models (chicken, pig, and cellular invasion), of which 21 genes belong to the flagellar system. This was in-line with our COG analysis, which revealed that motility-related genes were significantly overrepresented during host interaction (chicken, pig, and cellular invasion) (Fig 2 and Fig S5). Amongst the genes that were required across three host models and that did not belong to the flagellar system were *engD*, *livM* and *capM* (see Fig 3 and Fig 7). The EngD ortholog in *E. coli* (YchF) belongs to the GTPase family and is a negative regulator of the oxidative stress response [63]. YchF proteins have been implicated in pathogenesis of other bacterial species [63, 64]. With branched-chain amino acids (BCAA) being linked to chemotaxis [65], *livM*, part of the ABC-type BCAA transport system (*livM, livH, livK*, and *livJ*), may be implicated in chemotaxis within host organisms. Previous studies have shown a role for the *N*-linked protein glycosylation gene *capM* in chicken colonisation [41, 42], along with other members of the surface protein glycosylation locus, including *pglB, pglE* and *pglF* [66]. Our Tn-seq screen in the chicken colonisation model also identified Tn mutants in *pglB, pglF* and *pglI* exhibiting reduced colonisation (Table S3), whereas inactivation of *pglACD* resulted in reduced fitness during *in vitro* growth (Table S2). The PglH protein plays a crucial role in the glycan assembly process as it has a polymerase activity and adds the final *N*-acetlygalactosamine residues during surface decoration [67].

A total of 96 genes were only identified in the chicken colonisation screen and not during infection of gnotobiotic piglets, among these genes were the phosphate metabolism genes (*pstABS* and *phoR*). PstA, a gene that appears to be unrelated to motility (Fig S6), did not affect adhesion and invasion of human gut epithelial cells (Fig 7), and is vital for the capacity of *C. jejuni* to colonise chickens (Fig 3a,b). The expression of the phosphate regulon in *C. jejuni* 81-176 was increased in 1-day-old chickens, which suggests that phosphate levels might be low in chickens, thereby triggering the activation of the phosphate regulon [40]. The phosphate regulon might operate in a second messenger system that (in)directly regulates expression of chicken colonisation genes [68]. However, it remains to be investigated which *C. jejuni* genes are under control of this system. Tn mutants in genes involved in chemotaxis, *mcp4*_1 and *cheV*, were uniquely attenuated during chicken colonisation, whereas Tn mutants in *mcp4*_3 and *mcp4*_4 were uniquely attenuated during gnotobiotic piglet infection, suggesting the existence of host- or substrate-specific chemotaxis. A large number of genes involved in the utilisation of amino acids and organic acids such as lactate, pyruvate, acetate and tricarboxylic acid (TCA) cycle intermediates only appear to be required during gnotobiotic piglet infection. These host specific (central) metabolic requirements are most likely related to dietary differences between chickens and piglets. Metabolic pathways required across different hosts may be considered potential targets to develop antimicrobials. Our Tn library screens indicated that pyruvate kinase (*pyk*) is important in both chickens and piglets. Pyk is part of the Embden-Meyerhof-Parnas pathway, and shifting the metabolic flux at the level of Pyk has previously been suggested as a potential antimicrobial strategy [69].

A detailed analysis of the flagellar system across various *in vitro* and *in vivo* experimental models indicated that the extracellular part of the flagellum, the filament and hook structure, play a vital role in *C. jejuni* (Fig 8). This included genes encoding the filament chaperone gene FliW [16, 70] and the flagellar cap FliD. A FliD subunit vaccine has been shown to induce a transient but significant reduction (~2 log) of *C. jejuni* loads in the chicken caecum [71], demonstrating its potential as a target for intervention. Linked to the flagellar system is also the motility accessory factor 3 (*maf3*). We found that a *maf3* deletion mutant was motile, but displayed attenuated chicken colonisation and adhesion and invasion of Caco-2 cells. The *maf3* gene is located in the flagellar glycosylation locus [43]. However, the function of this gene is still unknown, and interestingly, isoelectric focusing analysis of a flagellar preparation from a *C. jejuni* 81-176 *maf3* deletion mutant did not indicate altered glycosylation of the flagellar filament [72].

Experimental models of *C. jejuni* survival and transmission, *i.e.* in the housefly and under low temperature conditions, were shown to be less stringent than the *in vivo* and *in vitro* host-pathogen models. This was reflected by subtler fold-changes compared to the host interaction models. This led to a lower number of candidate genes being identified and a lower conformation in validation experiments with defined gene deletion mutants (Fig 4 and Fig 5). Interestingly, this led to a novel potential target for intervention that is not linked to host interactions. The *trxC* gene was required for survival at low temperature, both in nutrient-rich and –poor conditions (Fig 5) and for resistance against peroxide (Fig 6), indicating a role in the oxidative stress response.

Although *C. jejuni* is considered the leading cause of bacterial gastroenteritis, our understanding of its biology is rather limited. Thus far, intervention strategies are unsuccessful in controlling *C. jejuni* in poultry and its transmission to humans. The development of future (novel) intervention strategies might best be aided by a thorough understanding of the biology of *C. jejuni* in its life cycle. The generated Tn mutant libraries in three well-characterised *C. jejuni* isolates represent a valuable tool for the *Campylobacter* research community. Our work in this study pointed out that many genes/pathways make indispensable contributions to the ability of *C. jejuni* to thrive in the host and environment. We anticipate that the use of the Tn mutant libraries in future studies will provide a continued insight into the mechanisms required for growth as well as survival within and outside host organisms. Perhaps most importantly, our comprehensive screening approach has again clearly shown that the flagella drive *C. jejuni* interaction with its hosts. Therefore, future efforts should focus on how to exploit this to effectively control infections caused by *C. jejuni*.

## Materials and Methods

### Ethics Statement

All chicken infection work was conducted in accordance with UK legislation governing experimental animals under project licence 40/3652 and was approved by the University of Liverpool ethical review process prior to the award of the licence.

### Bacterial Strains and Growth Conditions

Wild-type strains, defined gene deletion mutants, genetically complemented mutants and plasmids are summarised in Table S6. *C. jejuni* strains M1cam (derivative of M1 used in our laboratory) [16, 17], 11168 [10], and 81-176 [18] were cultured in Brain Heart Infusion (BHI) broth or on BHI agar supplemented with 5% (v/v) defibrinated horse blood in the presence of 5 μg/ml trimethoprim (TrM). Tn mutant libraries and defined gene deletion mutants were grown in the presence of 10 μg/ml chloramphenicol (Cm), whereas 50 μg/ml kanamycin (Km) was added to the media for culturing genetically complemented mutant strains. *C. jejuni* were grown under microaerophilic conditions (5% O2, 10% CO2, 85% N2) in a MACS VA500 Variable Atmosphere Work Station (Don Whitley, Shipley, United Kingdom). Liquid cultures were grown with agitation (200 rpm). For use in the experimental models, *C. jejuni* M1cam wild-type, defined gene deletion mutants and genetically complemented mutants were cultured for ~48 h, re-plated on fresh plates and grown for another 16 h. For Tn mutant libraries, freezer stocks were plated on 9 x 90 mm BHI blood agar plates (200 μl per plate) and grown for 16 h. *E. coli* NEB 5α or 10-β (New England Biolabs) were used for cloning and were cultured in Luria Bertani (LB) medium supplemented with appropriate antibiotics, at 37 °C.

### Construction of Defined Gene Deletion Mutants and Genetically Complemented Mutants

*C. jejuni* M1cam gene deletion mutants were constructed by allelic replacement of the gene with a chloramphenicol (*cat*) resistance cassette as described in de Vries *et al.* [16]. Gene deletion mutants were also subjected to phenotypic and genotypic characterisation, including motility (Fig S6) and *in vitro* growth in liquid culture (Fig S7). Whole genome sequencing (WGS) based variant analysis (single nucleotide polymorphisms [SNP] and insertion/deletions [INDELS]) of defined deletion mutants was used to screen for second-site mutations that might have affected the phenotypes under investigation. Variants were detected at 96 positions relative to the *C. jejuni* M1cam reference genome [16] (Table S5). The variant database was cross-referenced with data obtained in Tn mutant library screens to assess whether the gene affected by the variant had a potential impact on the phenotype under investigation. In addition, phenotypes of deletion mutants sharing a variant were compared to predict possible confounding effects of the second-site mutation. This in-depth analysis did not identify confounding effects of second-site mutations. For selected candidates, the respective defined gene deletion mutant was genetically complemented (Fig S8) and tested.

Genetic complementation of *mcp4*_1, *fliK*, *flaG*, *fliD*, *pstA*, *fabL*, *engD*, *livM*, *trxC*, *capM*, *maf3* and *flgK* mutants was performed using the pSV009 plasmid as described in de Vries *et al.* [16]. Predicted promoter region(s), including a ribosome-binding site (RBS) were derived from the Dugar *et al.* dataset [26]. For genes arranged in an operon, the promoter region was mostly identified upstream of the 5’ end of the transcript. In such cases, the expected promoter region was synthesised as a DNA string (GeneArt, Life Technologies, UK) flanked with appropriate restriction sites. Open reading frames (ORFs) of *fliK*, *fliD*, *pstA*, *engD* and *maf3* were cloned into *Xba*I and *BamH*I sites in pSV009, following which, respective promoter regions were inserted within the *Xho*I and *Xba*I sites, yielding the pSV009 derived plasmids containing the gene of interest fused to a suitable promoter. For *mcp4*_1, *flaG*, *fabL*, *trxC*, *capM* and *flgK*, a promoter consensus region was identified upstream of the start codon. A fragment containing the ORF plus ~200 bp upstream was cloned in the *Xho*I and *BamH*I sites of the plasmid pSV009. For *livM* no obvious promoter region could be located and therefore its transcription was placed under the control of a *cat* promoter by cloning the ORF into *Xba*I and *BamH*I sites of pSV009. Genetic complementation fragments, *i.e.* sequences for homologous recombination, a suitable promoter and ORF of the gene to be complemented, were amplified by PCR from the respective pSV009-derived plasmids using primers pSV009_GCampl_FW1/RV1. The genetic complementation fragments were introduced into respective M1cam defined gene deletion mutants by electroporation as described in de Vries *et al.* [16]. Oligonucleotide sequences are listed in Table S7.

### Gene Expression Analysis in Genetically Complemented Gene Deletion Mutants

For expression analysis of genes in complemented gene deletion mutants, total RNA was isolated as previously described in de Vries *et al.* [73]. DNA-free total RNA (500 ng) was reverse transcribed into cDNA using the QuantiTect reverse transcription kit (Qiagen). Real-time quantitative PCR (RT-qPCR) was performed using the SensiFast SYBR No-ROX Kit (Bioline) on a Rotor-Gene Q machine (Qiagen). Expression fold-changes relative to wild-type were calculated using the δδCt method [74] with the *gyrA* gene as a reference. For the isolation of total RNA, each bacterial strain was cultured in duplicate and RT-qPCR reactions were performed in duplicate.

### Analysis of Growth Kinetics

*C. jejuni* wild-type and gene deletion mutants were harvested from BHI blood agar plates in 2 ml of BHI broth. Culture suspensions were diluted to OD600nm ~0.2 and were used to analyse growth kinetics and motility. This suspension was used to inoculate 5 ml pre-warmed BHI broth at an OD600nm of ~0.005. Bacterial growth was monitored by optical density after 4, 8 and 24 h incubation at 200 rpm at 42°C under microaerophilic conditions (*n* ≥ 3). Ten-fold serial dilutions were made and plated onto BHI blood agar plates.

### Analysis of Motility

Motility assays (*n* = 3) were performed in semi-solid agar as described in de Vries *et al.*[16]. Bacterial suspensions that were diluted to OD600nm ~0.2 (see previous section) were used to stab 90 mm petri dishes containing 25 ml of BHI broth supplemented with 0.4% (w/v) agar. These plates were incubated under microaerophilic conditions at 42°C for 24 h before measuring the diameter of growth/motility.

### Genome Sequencing of Defined Gene Deletion Mutants

Libraries for Illumina sequencing were prepared using the NEBNext Ultra or Ultra II DNA library prep kit (New England Biolabs) and sequenced on the MiSeq platform as described in de Vries *et al.* [16]. For variant analysis, reads were mapped to the M1cam reference genome (accession no. CP012149 [16]) using Stampy [75], variants were identified with Samtools [76] and the effect at the protein level was predicted using SnpEff [77]. Gene deletion mutants in *pflA* and *flaD* were analysed previously in de Vries *et al.*[16].

### Construction of Tn Mutant Libraries in *C. jejuni*

A Tn donor plasmid suitable for Tn-seq [19] was constructed by amplifying the *mariner* Tn encoding the Cm-resistance cassette from pAJG39 [36] using a single 5'- phosphorylated primer PBGSF20. This introduced *Mme*I restriction sites within the inverted repeats of the Tn element. The Tn element was sub-cloned into pJET1.2 (Thermo Scientific) and the resulting plasmid pSV006, was used for *in vitro* Tn mutagenesis as described in Holt *et al.*, [78] with minor adjustments. Briefly, 2 μg of *C. jejuni* DNA was incubated for 5 h at 30°C with 1 μg of pSV006 and ~250 ng Himar1-C9 transposase, which was purified as described in Akerley *et al.* [79]. After end-repair with T4 DNA polymerase and *E. coli* DNA ligase the mutagenesised DNA was transferred to *C. jejuni* by natural transformation [16]. Tn transformants were harvested from plates and pooled. The pooled library was used to inoculate 100-200 ml BHI-TrM-Cm broth to an OD600nm of ~0.1 and grown overnight). This yielded the “working stocks” that were used in further experiments. Chromosomal DNA was isolated with Genomic-tip columns (Qiagen).

### Gene Fitness Analysis

Tn insertion sites were identified using Tn-seq [19], essentially as described in Burghout *et al.*, [80]. Briefly, Tn mutant library DNA was fragmented by *Mme*I restriction digestion. Adapters containing inline barcodes were ligated to the *Mme*I fragments, and amplified using the primers PBGSF29 and PBGSF30 with NEBNext high fidelity polymerase (New England Biolabs). Tn insertion sites were sequenced using single-end 40-50 bp sequencing on the Illumina HiSeq 2500 platform. Sequence reads were demultiplexed using the FastX toolkit barcode splitter and analysed further with the ESSENTIALS pipeline [22]. Sequence reads were aligned to the *C. jejuni* genomes [16, 18, 81] with a match of ≥16 nt. Kernel density plots were generated in R to distinguish “true” Tn insertions from “noise” sequencing reads, yielding a read count cut-off per Tn library. Insertion “hot spotting” was analysed by plotting the Log2 read count per chromosomal position using an in house Perl script. As a measure of gene fitness, the Log2 fold-change of observed *vs* expected reads was calculated per gene, with Kernel density plots allowing accurate delineation of fitness (required for *in vitro* growth) and non-fitness genes [22]. Additional criteria were: a Benjamini & Hochberg adjusted *P* < 0.05 and a probability that the gene was inactivated by a Tn insertion of > 0.95, as calculated using a derivative of Poisson’s law; 1-e^N x Ln(1-f)^, with N = number of unique Tn insertion mutants and f being the gene size divided by the size of the genome. In addition, genes for which no sequence reads were detected and the probability of inactivation was > 0.95 were considered to be required for fitness. For functional class enrichment analysis COGs were assigned to M1cam, 11168 and 81-176 proteins and consensus COGs were assigned to homologous groups (HGs; see next section). The overrepresentation of COG classes was assessed using a Fisher exact test with *Q*-value multiple testing correction [82].

### Identification of Homologs in *C. jejuni* Strains

Protein sequences of *C. jejuni* M1cam, 11168 and 81-176 were clustered into putative homologous groups (HGs) with OrthAgogue [83]. For this, a collective database was generated and proteins of the three strains were queried against this database. The reciprocal best-hit protein pairs were identified by applying an e-value filter cut-off of 1e-5 to the “all against all” BLAST output (only proteins > 50 amino acids in length were included). The putative HGs were identified by clustering with MCL by applying different values of an inflation parameter that defines cluster tightness [84]. After testing different inflation parameters, an inflation value of 2.6 was selected. This ensured that the majority of core HGs had only one representative gene, keeping the number of HGs with duplicated genes from an isolate to a minimum.

### Systematic Literature Review on *C. jejuni* 11168 Defined Gene Deletion and Tn Mutants

To assess which *C. jejuni* genes have been deleted in strain 11168, a PubMed search was conducted in July 2015 with the following search terms: “*Campylobacter jejuni*” and “mutant”. Publications were screened for the description of defined gene deletion mutants.

### Chicken Colonisation Experiments

One-day-old Ross 308 broiler chicks were obtained from a commercial hatchery. Chicks were housed in the University of Liverpool, High Biosecurity Poultry unit. Chicks were individually tagged with leg rings or wing bands and maintained in floor pens at UK legislation recommended stocking levels allowing a floor space of 2,000 cm2 per bird at 25°C on wood-shavings litter that was changed weekly prior to inoculation, and were given *ad libitum* access to water and a pelleted laboratory grade vegetable protein-based diet (SDS). Prior to experimental infection, all birds were confirmed as *Campylobacter*-free by taking cloacal swabs, which were streaked onto selective blood-free agar (mCCDA) (Lab M) supplemented with *Campylobacter* Enrichment Supplement (SV59) and grown for 48 h at 42°C under microaerophilic conditions.

At 21 days of age, birds were inoculated by oral gavage with ~1.8 x 10^8^ CFU *C. jejuni* M1cam Tn library ‘C’. The birds were split into five cages (*n =* 6, 7, 7, 6, 6; group 1-5, respectively). Six days post-inoculation (p.i.), birds were killed by cervical dislocation. At necroscopy the caeca were removed aseptically and caecal content was collected. Next, the caecal contents were diluted 5- and 50-fold with Maximum Recovery Diluent (MRD) and 0.5 ml was plated per mCCDA-Cm (10 plates in total per dilution) for recovery of Tn mutants that successfully colonised the birds. Ten-fold dilution series were plated on mCCDA-Cm for enumeration. After 2 days growth, *C. jejuni* M1cam Tn mutants were recovered from the plates by scraping in 2 ml MDR and pelleted by centrifugation. The resulting pellets were stored at -80°C. DNA was isolated from plate harvest pellets with Genomic-tip columns (Qiagen).

The composition of the input Tn mutant library (*n* = 3) and the library recovered per cage of birds (*n* = 4, cage 1-4), a cage was considered as a single unit of colonisation, was analysed by Tn-seq [19]. For this, 1 μg of DNA was pooled per group of birds (*n* = 6, 7, 7, 5 for groups 1 to 4, respectively). One bird from group 4 and 2 birds from group 5 could not be analysed due to heavy contamination on the recovery plates. As a result of this group 5 was eliminated from further analysis.

For validation with *C. jejuni* M1cam defined gene deletion mutants and genetically complemented mutants, birds were inoculated with ~1.8 x 108 CFU, at 6 days p.i. chickens were killed by cervical dislocation, the caeca were removed aseptically and the caecal contents plates onto mCCDA plates for enumeration.

### Infection of Houseflies

Newly pupated female houseflies (*Musca domestica*) were inoculated with 1 μl *C. jejuni* suspension *via* their proboscis [85]. For screening of Tn mutant survival, 5 groups of 10 flies were inoculated with ~10^6^ CFU M1cam Tn library ‘C’ on four different days. The flies were incubated for 4 h at 20°C (in the dark) after which flies were homogenized using a Drigalsky spatula and 5 ml BHI was added. Large debris was removed through a low speed (700 x *g*) spin and the supernatant was adjusted to 5 ml with BHI before plating 0.5 ml per mCCDA-Cm (10 plates in total). Ten-fold dilution series were plated on mCCDA-Cm for enumeration. After 24 h growth, Tn mutants were recovered from plates by scraping in 2 ml BHI medium, centrifuged and pellets were stored at -80°C for DNA isolation with Genomic-tip columns (Qiagen). Chromosomal DNA from the 5 groups of 10 flies per day was pooled; resulting in a single pooled sample for the library recovered in each of the 4 replicate experiments. The composition of the input Tn mutant library (*n* = 4) and the library recovered per group of flies (*n* = 4) was analysed by Tn-seq [19].

For validation experiments with *C. jejuni* M1cam gene deletion mutants (*n* ≥ 4) and genetically complemented mutants (*n* = 6), groups of 5 flies were inoculated with ~106 CFU, killed 4 h p.i., and the bacterial load was quantified from pools of five flies or five flies individually (only for 2 replicates with gene deletion mutants), in both cases the average CFU per fly was used to calculate the Log10 decrease of CFU per fly relative to the inoculum.

### Cold Survival Assay

A suspension was prepared from *C. jejuni* M1cam Tn mutant library ‘C’ plate harvests, to an OD_600nm_ ~ 0.5 (~5 x 108 CFU/ml). The Tn mutant library suspension was harvested by centrifugation and resuspended in either BHI or sterile tissue-culture grade water (experiment 1) or chicken juice, tap water or rain water (experiment 2). The chicken juice was prepared as follows: 10 frozen whole chickens were purchased from a commercial supplier and were allowed to defrost within the packaging for at least 16 h, as described by Brown *et al.[86].* Concentrated liquid was recovered (> 200 ml), sterilised through a 0.2 μM filter and stored at–20 °C. Tap water was obtained from a mains-fed tap in our laboratory that was allowed to run for at least 2 min prior to collecting the water used in experiments. To collect rain water, a large tub was left outside our laboratory overnight on a rainy evening. The rain water and tap water were passed through a 0.22 μM filter unit and stored at 4°C. Chicken juice, tap water and rain water were also screened for the presence of *Campylobacter* spp. by plating on mCCDA plates. The Tn library samples were incubated at 4°C and aliquots were taken at 0 h, 6 h, 1 and 7 days for experiment 1 (*n* = 3) or at 1, 3, and 7 days for experiment 2 (*n* = 4) and plated for recovery of Tn mutants. Ten-fold dilution series were plated on BHI-TrM-Cm for enumeration. After 2 days incubation, Tn mutants were recovered from plates by scraping in 2 ml BHI per plate, centrifuged and pellets were stored at -80°C for DNA isolation with Genomic-tip columns (Qiagen). The Tn library composition at time = 0 h was compared to the library recovered after incubation at 4°C under the above mentioned conditions by Tn-seq [19].

For validation experiments, the survival of gene deletion mutants and genetically complemented mutants was assayed as described above (*n* = 4).

### Infection of Human Gut Epithelial Cells

Caco-2 cells (ATCC CC-L244 HTB-37), were cultured in DMEM (Life Technologies) supplemented with 10% (v/v) heat inactivated FBS (Gibco), and 1% (v/v) non-essential amino acids (Sigma Aldrich), at 37 °C with 5% CO_2_.

The *C. jejuni* M1cam Tn mutant library ‘C’ was used to infect seven 143 cm^2^ dishes per replicate (*n* = 4) containing a monolayer of Caco-2 cells at a multiplicity of infection (MOI) of 100 in low phosphate HEPES buffer (10 mM HEPES, 5.4 mM KCl, 145 mM NaCl, 5 mM glucose, 1 mM MgCl2, 1 mM CaCl2, 2 mM phosphate buffer pH 7.4) [87]. Cells were incubated for 2 h after which the non-adherent fraction from two dishes was recovered onto BHI blood agar plates. For adherence, two dishes were washed three times in Dulbecco’s PBS (D-PBS), Caco-2 cells were lysed in 10% (v/v) Triton-X100 solution in D-PBS, and bacteria were recovered on BHI blood agar plates. The remaining five dishes were washed three times in D-PBS and incubated for an additional 2 h in DMEM with 250 μg/ml gentamycin. Tn mutants that invaded Caco-2 cells were recovered after washing three times in D-PBS and Caco-2 cell lysis in 10% (v/v) Triton-X100 in D-PBS. After two days of growth on plates, Tn mutants were recovered from the plates in 2 ml BHI, centrifuged and pellets were stored at -80°C. DNA was isolated from harvested pellets using Genomic-tip columns (Qiagen).

Validation of the screen was performed using 24-well plates, for which Caco-2 cells were infected with various *C. jejuni* M1cam defined gene deletion mutants and genetically complemented strains (*n* ≥ 4) as described above. Bacteria recovered from different fractions were serially diluted and plated on BHI agar plates containing appropriate antibiotics. Adhesion and invasion of Caco-2 cells by various *C. jejuni* strains was calculated relative to the matched wild-type (*n* ≥ 3).

To test the effect of competition, Caco-2 cells were infected at an MOI of 100 with a mix of the wild-type strain to a defined gene deletion mutant at a ratio of 100:1. A competitive index (CI) score was calculated by dividing the ratio of mutant to wild-type recovered from adherent and invaded fractions by the ratio of mutant to wild-type bacteria in the inoculum (*n* ≥ 4).

### Analysis of Conditionally Essential Genes

Tn-seq data from the different experimental models (conditionally essential genes screens) was processed as described in “gene fitness analysis”. To identify genes of which Tn mutants were attenuated or enriched in the tested models, read counts were collected per gene and compared between output (recovered) and the input or control conditions (as defined above). Only genes covered by > 100 reads in the control condition were considered, allowing assessment of 809 ± 58 (67 ± 5%) non-fitness genes (define ‘non-fitness genes’) in the selected models. The following filter steps were applied: a Log2 fold-change (FC) below the attenuated cut-off value or higher than the enriched cut-off value, Benjamini & Hochberg false discovery rate < 0.05, and two or more Tn mutants showing a Log2 fold-change below the attenuated cut-off value or higher than the enriched cut-off value (analysed using a custom Python script). The Log2 fold-change cut-offs were selected based on MA-plots. In addition, the 514 genes that were obligate essential or required for fitness (defined above) were eliminated from the analysis, see “gene fitness analysis”. COG functional class enrichment was analysed using a Fisher exact test with *Q*-value multiple testing correction [82].

### Hydrogen Peroxide Sensitivity Assay

*C. jejuni* strains were added to pre-cooled BHI agar (~45°C) to a calculated OD600nm ~0.005 and 25 ml of this media-bacterial suspension was poured into 90 mm petri dishes. Blank filter paper discs, 6 mm, were loaded with 10 μl of 0.05, 0.1, 0.25, 0.5, 1, 2.5, 5 or 7.5 M H2O2 solution. The discs were allowed to air dry and were then placed in the center of the solidified agar plates. The plates were incubated for 24 h, after which time the inhibition zone diameter was measured using a ruler (*n* = 4).

### Sequencing Data

Tn-seq and genome sequencing data has been deposited in the European Nucleotide Archive (http://www.ebi.ac.uk/ena) and are available *via* study accession number [This will be openly available after acceptance and can be made available for review].

### Statistical Analysis

Statistical analysis was performed in GraphPad Prism v6.

## Acknowledgements

We thank Gemma Murray from the Department of Genetics, University of Cambridge for providing the R script used for COG assignment and Diane Newell for providing the *C. jejuni* M1 strain. We thank David Lampe for providing the pMALC9 plasmid and Roy Chaudhuri for sharing the read count mapping Perl script.

## Author contributions

Conceptualization: NW, PW, TH, DJM, AJG. Methodology: SPWdV, SG, AB, EW, AW, ANJ, LLL, SH, HK, KM, PE, NW, PW, AJG. Software: AB, FMM. Validation: SPWdV, SG, AB, AJG. Formal analysis: SPWdV, SG, AB, ANJ, AJG. Investigation: SPWdV, SG, EW, AW, ANJ, LLL, SH, SK, KM, EP, DPW, JL'H, DGES, PE, NW, PW. Resources: FMM, AZ. Data curation: SPWdV, SG, AB. Writing original draft: SPWdV, SG, AG. Writing review and editing: SPWdV, SG, DJM, AG. Visulaization: SPWdV, SG, AG. Supervision: AJG. Project administration: AJG. Funding acquisition: TH, DJM.

## Supporting Information

**Table S1. Overview of Tn mutant libraries constructed in *C. jejuni* M1cam, 11168, and 81-176.**

**Table S2. Overview gene fitness analysis in *C. jejuni* M1, 11168 and 81-176.** As a measure of gene fitness, the Log2 fold-change of observed *vs* expected reads was calculated per gene, with Kernel density plots allowing accurate delineation of fitness and non-fitness genes in each *C. jejuni* strain. Additional selection criteria were: a Benjamini & Hochberg adjusted *P* < 0.05 and a probability that the gene was inactivated by a Tn insertion of > 0.95, as calculated using a derivative of Poisson’s law. In addition, genes for which no sequence reads were detected and the probability of inactivation was > 0.95 were included. For comparative analysis of fitness genes in the three *C. jejuni* strains, homologs were identified; see Materials and Methods for a detailed description.

**Table S3. Overview of conditionally essential gene analysis.** To identify genes of which Tn mutants were attenuated or enriched in the tested experimental models, read counts were collected per gene and compared between output (recovered) and the input or control conditions. The following filter steps were applied: genes represented > 100 reads in the input/control condition, a Log2 fold-change (FC) below the attenuated cut-off value or higher than the enriched cut-off value, Benjamini & Hochberg false discovery rate < 0.05, and two or more Tn mutants showing a Log2 fold-change below the attenuated cut-off value or higher than the enriched cut-off value. The Log2 fold-change cut-offs were selected based on MA-plots. In addition, the 514 genes that were obligate essential or required for fitness in *C. jejuni* M1cam (Table S2) were eliminated from this analysis.

**Table S4. Conditionally essential genes grouped per functional (COG) category**. Effect of Tn insertions on the ability of *C. jejun*i M1cam to colonize commercial broiler chickens, infect gnotobiotic piglets, adhere and invade human gut epithelial tissue culture cells, survive in houseflies and at 4°C in various media (chicken juice, BHI, tap water, rain water, and sterile water). Genes of which Tn mutants showed significantly attenuated or enriched fitness in the tested experimental models are listed and are grouped according to their COG functional classification. Data represented as Log2 fold-change is also presented in Fig 2a. Orange = significantly attenuated, Blue = significantly enriched, and Grey = no significant Tn-seq Log2 fold-change.

**Table S5. Whole genome sequencing (WGS) based variant analysis of *C. jejuni* M1cam defined gene deletion mutants.** For variant analysis, reads were mapped to the *C. jejuni* M1cam reference genome [16] with Stampy, variants (single nucleotide polymorphisms [SNP] and insertion/deletions [INDELS]) detected using Samtools, and the effect at the protein level was predicted using SnpEff. Cross-referencing the variants with data obtained in the Tn mutant library screens as well as phenotypes of defined gene deletion mutants sharing a variant did not reveal any confounding effects of second-site mutations, suggesting that the observed phenotypes for gene deletion mutants were true.

**Table S6. Bacterial strains and plasmids used in this work.**

**Table S7. Oligonucleotides used in this study.**

**Fig S1**. **Distribution of Tn insertion sites in *C. jejuni* M1cam, 11168 and 81-176 Tn mutant libraries.** Plotted are Log2 reads per Tn insertion above the read count cut-off (see Table S1 and Materials and Methods).

**Fig S2. Overlap of the Tn insertions per mutant library in *C. jejuni* M1cam, 11168, and 81-176 Tn libraries**. Only Tn insertion sites above the read count cut-off are included in this analysis (see Table S1 and Materials and Methods).

**Fig S3. Circular genome visualisation indicating fitness genes in *C. jejuni* M1cam, 11168 and 81-176.** Genes required for fitness in one, two or all three of the strains are coloured-coded accordingly (See Table S2). A detailed description of the selection criteria is provided in the Materials and Methods section.

**Fig S4. Complexity of the *C. jejuni* M1cam Tn mutant library ‘C’ (Table S1) in the conditionally essential gene screens as analysed by Tn-seq.** Tn mutants represented by > 10 reads were considered to be present. Of note, in the “housefly survival” and “survival under nutrient-rich/poor conditions” screens, a higher number of Tn mutants were detected in the recovered (output) samples, which was the result of a lower sequence depth for some replicates of the input/inocula. Data is shown as individual data points and bars representing the mean with SEM. Statistical significance was analysed using a Mann-Whitney test with * *P* < 0.05 and ** *P* < 0.01.

**Fig S5. Functional class (COG) enrichment analysis of genes required during colonisation of chickens, infection of gnotobiotic piglets, invasion of human gut epithelial tissue culture cells, or survival in houseflies.** The overrepresentation of COG classes was assessed using a Fisher exact test with *Q*- value multiple testing correction; * *Q* < 0.05 and *** *Q* < 0.001.

**Fig S6. Motility of *C. jejuni* M1cam defined gene deletion mutants**. The data is represented as percentage relative to the wild-type (*n* ≥ 4). Statistical significance was calculated using Kruskal-Wallis with Dunn’s correction for multiple comparisons with * *P* < 0.05, ** *P* <0.01 and *** *P* < 0.001.

**Fig S7. Growth kinetics of *C. jejuni* M1cam defined gene deletion mutants.** Growth of *C. jejuni* M1cam wild-type and gene deletion mutants were recorded as viable counts over 24 h at 42°C under microaerophilic conditions. Data is represented as mean and SD (*n* ≥ 2).

**Fig S8. Genetic complementation of *C. jejuni* M1cam defined gene deletion mutants.** RT-qPCR analysis revealed the restored expression of *mcp4_1*, *flaG*, *pstA*, *fabL*, *livM*, *fliW*, *maf3* and *flgK* in genetically complemented stains. Lower expression of *fliD*, *trxC* and *capM* was observed in respective genetically complemented strains, compared to the wild-type. Although the *fliD* gene was expressed in the genetically complemented mutant, albeit at lower levels than the wild-type, it was unable to restore motility (data not shown). This might be (partially) caused by deregulated flagellar assembly due to an increased expression (2.9-fold compared to wild-type) of *fliS* (a flagellar chaperone [88]) located downstream of *fliD* (data not shown). WGS analysis did not reveal any genomic variations in the *fliD* mutant linked to motility (Table S5). The data is represented as Log2 fold-change relative to the expression levels of the candidate gene in wild-type. Statistical significance (n ≥ 4) of mutant *vs* wild-type and genetically complemented strains *vs* wild-type was calculated using a one-way ANOVA test with Bonferroni correction for multiple comparisons. For all experiments, data shown are means and SD. * *P* < 0.05, *** *P* < 0.001

